# CofActor: A light and redox-gated optogenetic clustering tool to study abnormal cytoskeletal dynamics in stressed cells

**DOI:** 10.1101/857706

**Authors:** Fatema B. Salem, V. V. Prabhu, Abu-Bakarr Kuyateh, Wyatt Paul Bunner, Alexander Murashov, Erzsebet M. Szatmari, Robert M. Hughes

## Abstract

The biochemical hallmarks of neurodegenerative diseases (neural fibrils, clumps, and tangles; heightened reactive oxygen species (ROS); cofilin-actin rods) have presented numerous challenges for development of *in vivo* diagnostic tools (1–7). Biomarkers such as amyloid beta (Aβ) fibrils and Tau tangles in Alzheimer’s Disease (AD) are only accessible via invasive cerebrospinal fluid assay for peptide mass fingerprinting or post-mortem diagnosis (8–11), while ROS can be fleeting and thus challenging to monitor *in vivo* (12–15). While remaining a challenge for *in vivo* detection, the unique protein-protein interactions underlying these disease-specific biomarkers also present rich opportunities for the engineering of *in vitro* pathology-sensitive biosensors and bioactuators. These tools can be useful for the investigation of critical, early stage events in neurodegenerative diseases in both cellular and animal models (16, 17), while potentially leading to advanced detection reagents with clinical applications. In this report, we describe a light and redox-gated protein switch inspired by the phenomenon of cofilin-actin rod formation, which occurs in stressed neurons in the AD brain and following brain ischemia (18). By coupling the redox-sensitive interaction of cofilin and actin with the light responsive Cry2-CIB blue light switch, we accomplish both light- and ROS-gated control of this interaction. The resulting switch is referred to as the “CofActor” system. Site-directed mutagenesis of both cofilin and actin partners demonstrate which residues are critical for sustaining or abrogating the light and redox gated response. Furthermore, switch response varies depending on whether oxidative stress is generated via glycolytic inhibition or a combination of glycolytic inhibition and azide-induced oxidative stress. Finally, light and redox gated switch function was demonstrated in cultured hippocampal neurons. As a dual input biosensor, CofActor holds promise for the tracking of early stage events in neurodegeneration and the investigation of actin-binding protein interactions under oxidative stress.

## INTRODUCTION

Actin-cofilin bundles or ‘rods’, formed in neurons as part of an oxidative stress response, impact synaptic function and may play a significant early role in the progression of neurodegenerative disorders (19). Prior studies have shown that actin and cofilin combine in a 1:1 ratio in rods, with actin assuming a highly twisted conformation (20). This unusual arrangement excludes most other potential binding partners, with particular isoforms of 14-3-3 protein being a notable exception (20, 21). While cofilin-actin rods are frequently observed in post-mortem brain slices from AD patients, their transient nature, along with the multi-faceted activities of both cofilin and actin, makes them particularly challenging to study *in vivo* (19). Thus, methods for the induction of cofilin-actin rods in cell culture have been developed, including in primary neurons exposed to Aβ fibrils (22). In addition, application of an ATP-depletion cocktail comprised of sodium azide and D-deoxyglucose is a potent promoter of cofilin-actin rod formation in HeLa cells (20).

Earlier work shows that mutagenesis of only one or two amino acids in cofilin (e.g. S3E; S3A.S120A) can significantly reduce the affinity of cofilin for native actin structures, both *in vivo* and *in vitro* (23–26). These mutant cofilins are subsequently able to function as biosensors and bioactuators of cofilin-dependent cell functions (27–29). Recently, a cofilin mutant (R21Q) was described that enables real-time tracking of cofilin-actin rod formation and dissipation in cell culture models when expressed as a fusion with the fluorescent protein mCherry (30). Unlike overexpressed wild type cofilin, the R21Q mutant resists incorporation into cofilin-actin rods in the absence of oxidative and energetic stress; however, it readily incorporates into cofilin-actin bundles under oxidative and energetic stress. In light of the R21Q study, we asked whether a cofilin mutant with substantially reduced affinity for F-actin could be the basis for an optogenetic switch that enables the bundling of cofilin and actin in response to both light and redox stimuli.

The requirement for such a switch would be that it resists background incorporation into cofilin-actin bundles under both normal and oxidative stress conditions, yet readily forms cofilin-actin bundles when exposed to a combination of oxidative stress and light. However, in contrast to native cofilin-actin bundle formation, reversal of the light induced cofilin-actin bundles could be accomplished without removal of the source of redox stress. With the advantage of rapid & reversible light mediated induction, the resulting switch would allow for tracking the formation and movement of cofilin-actin bundles in living cells, including neurons.

## MATERIALS AND METHODS

### Plasmids and Cloning

Cloning of Cof.Cry2PHR.mCh and Cry2PHR.mCh.Cof constructs has been previously reported (31). Cloning of the β-Actin.Cib.GFP construct was conducted according to previously reported methods (32). Briefly, genes encoding β-Actin and Cib were PCR amplified with overlapping primer sequences. Gene fragments were gel isolated, then stitched together via splice overlap extension (SOE) PCR. The resulting gene fragment was trimmed via restriction digest, followed by ligation into complementary restriction sites in the target plasmid. Point mutations were introduced into Actin in a pNic28 plasmid, followed by subcloning into the Cib.GFP mammalian expression plasmid (phCMV-GFP;Genelantis). Cofilin mutants were synthesized by IDT, PCR amplified, and subcloned into the Cry2.mCh mammalian expression plasmids (mCherry-N1 (Clontech)) via restriction digest and ligation. The actin encoding gene (pCAG-mGFP-actin; Addgene #21948) construct was a generous gift from Ryohei Yasuda (33).

### Cell lines and transfection

Midi prep quantities of DNA of each construct were created from *E. coli* and collected for cell transfection. Transfection of HeLa cells was then performed with the Lipofectamine 3000 reagent (Invitrogen) following manufacturer’s suggested protocols. Briefly, for dual transfections in 35 mm glass bottom dishes, plasmid DNA was combined in a 1:1 ratio (1,250 ng per plasmid) in 125 ul of Opti-Mem, followed by the addition of 5 ul of P3000 reagent. For single transfections in 35 mm glass bottom dishes, 2,500 ng of plasmid DNA was used per transfection. In a separate vial, 3.75 ul of Lipofectamine 3000 were added to 125 ul of Opti-Mem. The two 125 ul solutions were combined and allowed to incubate at room temperature for 10 min, followed by dropwise addition to cell culture. Transfection solutions were allowed to remain on cells overnight. Cells were maintained at 37°C and 5% CO2 in a humidified tissue culture incubator, in culture medium consisting of DMEM supplemented with 10% FBS and 1% Penicillin-Streptomycin.

### ATP Depletion and Immunofluorescent staining

#### Fixed cell experiments

Transfected HeLa cells were washed with Dulbecco’s PBS (with calcium and magnesium; 3 × 1 mL), prior to treatment with ATP depletion medium (6 mM D-Deoxyglucose and 10 mM Sodium Azide in Dulbecco’s PBS) for the indicated time intervals. ATP depletion medium was removed by aspiration, cells washed gently (1X with 1 mL Dulbecco’s PBS), then fixed for 45 min with pre-warmed 4% Paraformaldehyde solution (37°C; prepared from 16% PFA (Electron Microscopy Sciences and DPBS) at room temperature for 45 min. Following fixation, cells were washed with PBS, then permeabilized for 3 min using pre-chilled methanol (−20°C). Methanol was removed by aspiration, then cells were blocked for 30 min with antibody dilution buffer (30 µl Triton X-100, 0.1 g of BSA, 10 mL of Dulbecco’s PBS). Cells then were incubated overnight at 4°C with primary antibody (anti-Actin (Santa Cruz); 1:500 in antibody dilution buffer). The following day, primary antibody solution was removed by pipette, and cells were washed three times with Dulbecco’s PBS. Cells were incubated with Alexa 488 conjugated goat anti-mouse secondary (Invitrogen; 1:200 in antibody dilution buffer) for 1 hour at room temperature, followed by a Dulbecco’s PBS wash (1 mL; 3 × 5 min). Cells were stored in Dulbecco’s PBS prior to imaging.

##### Live cell experiments

Transfected HeLa cells were washed Dulbecco’s PBS (with calcium and magnesium; 1 × 1 mL), prior to treatment with ATP depletion medium (6 mM D-Deoxyglucose and 10 mM Sodium Azide in PBS). Cells were allowed to equilibrate in the live cell incubation system (OKOLab) for 10 minutes prior to beginning the illumination sequence.

### Imaging

#### Confocal Microscopy

Confocal images of fixed cells were obtained with Olympus IX2-DSU tandem spinning disk confocal laser scanning microscope or with a Zeiss LSM 880 microscope with Airyscan technology. Fluorescence images were colorized and overlaid using FIJI software.

#### Widefield Microscopy

A Leica DMi8 Live Cell Imaging System, equipped with an OKOLab stage-top live cell incubation system, LASX software, Leica HCX PL APO 63x/1.40-0.60na oil objective, Lumencor LED light engine, CTRadvanced+ power supply, and a Leica DFC900 GT camera, was used to acquire images. Exposure times were set at 150 ms (GFP, 470 nm) and 300 ms (mCherry, 550 nm), with LED light sources at 50% power, and images acquired every 30 seconds over a 20 min time course.

### Primary neuron culture preparation, transfection and treatment

Dissociated hippocampal neuron cultures were prepared as previously described (34). Briefly, hippocampi from newborn B6 mice were used to prepare dissociated postnatal neuron cultures. Cells were plated into 35 mm glass bottom Petri dishes with growth medium consisting of BME supplemented with 10% BCS and antibiotics. 24 h after plating, the culture medium was changed to Neurobasal A medium, supplemented with B27-plus reagent (Invitrogen). Neurons were transfected with the CofActor optogenetic system (6 µg plasmid/plate) on day in vitro 3-5 (DIV3-5) using Lipofectamine 2000 (Invitrogen). 48 hours post transfection, culture medium was removed, and neurons were placed in ATP depletion medium (6 mM D-Deoxyglucose and 10 mM Sodium Azide in PBS). Cultures were returned to tissue culture incubator (37°C; 5% CO_2_) and allowed to equilibrate for 10 minutes prior starting the illumination sequence. Live cell imaging of neurons was performed before and after illumination, using a Keyence (BZ-X800) all-in-one fluorescence microscope.

### Particle counting

Analysis of imaging data was performed in FIJI equipped with the BioFormats package. Particle counting was performed using the Analyze Particles feature. Particle size was restricted from 20-200 pixels and circularity restricted from 0.2 – 1.00. Particle counts are reported as average number of particles counted per cell.

### Western blotting

HeLa cells were lysed post-transfection with 200 µL of M-PER lysis buffer (Thermo Scientific) plus protease inhibitors. After 10 min on a rotary shaker at room temperature, lysates were collected and centrifuged for 15 min (94 rcf; 4°C). The supernatants were used for further experiments. The resulting lysates were subjected to electrophoresis on a 10% SDS-PAGE gel and then transferred onto PVDF membranes (20 V, overnight, at 4°C). Membranes were then blocked for 1 h with 5% BSA in TBS with 1% Tween (TBST), followed by incubation with primary antibody (Anti-mCherry antibody (Cell Signaling); 1:1000 dilution in 5% BSA – TBST; Anti-GFP antibody (Santa Cruz); 1:1000 dilution in 5% BSA – TBST) overnight at 4°C on a platform rocker. The membranes were then washed 3 × 5 min each with TBST and incubated with the appropriate secondary antibody in 5% BSA – TBST for 2 hours at room temp. After washing 3 × 5 min with TBST, the membranes were exposed to a chemiluminescent substrate for 5 min and imaged with using an Azure cSeries imaging station.

## RESULTS

Prior studies demonstrated that point mutants of Cofilin-1 (e.g. S3E, S3A.S120A) exhibit diminished actin binding capabilities, both when expressed as standalone cofilins and when incorporated into protein fusions (Cofilin – Cryptochrome 2 – mCherry) (23). We investigated whether the same cofilin-containing protein fusions exhibited a similar response to oxidative and energetic stress in the presence of an ATP depletion media, as has been previously reported for both endogenous and overexpressed cofilins (20, 30, 35). Following literature procedures, cofilin (WT, S3E, S3A, S3A.S120A)-Cry2PHR-mCh protein fusions expressed in HeLa cells were incubated with either PBS or ATP depletion media for 1 h, followed by fixation and imaging. As a result of the applied oxidative stress, cofilins with full actin-binding capability (WT and S3A) were readily incorporated into clusters, whereas cofilins with impaired actin-binding ability (S3A.S120A and S3E) failed to incorporate into clusters (Figure 1). Via immunostaining with an actin antibody, we demonstrated that the observed cofilin clusters were co-localized with endogenous β-actin (Fig. 1B). Further exploration of the ATP depletion buffer revealed that using the full complement of redox and energetic stress (10 mM NaN3/6 mM D-deoxyglucose) resulted in abundant cofilin-actin clusters distributed throughout the cell body, whereas treatment with glycolytic inhibition only (0 mM NaN3/6 mM D-deoxyglucose) resulted in the accumulation of cofilin-actin bundles around the cell periphery (Figure 2 and Supporting Figures 1-3). We attribute this effect to naturally occurring ROS production, which is higher at the highly dynamic cell edges (36). Experiments were also conducted to establish the minimum time and concentrations necessary for induction of cofilin-actin clustering, demonstrating that 15 minutes in full ATP depletion medium was sufficient for inducing robust cofilin-actin bundling (Supp. Fig. 1 – 3).

**FIGURE 1.**
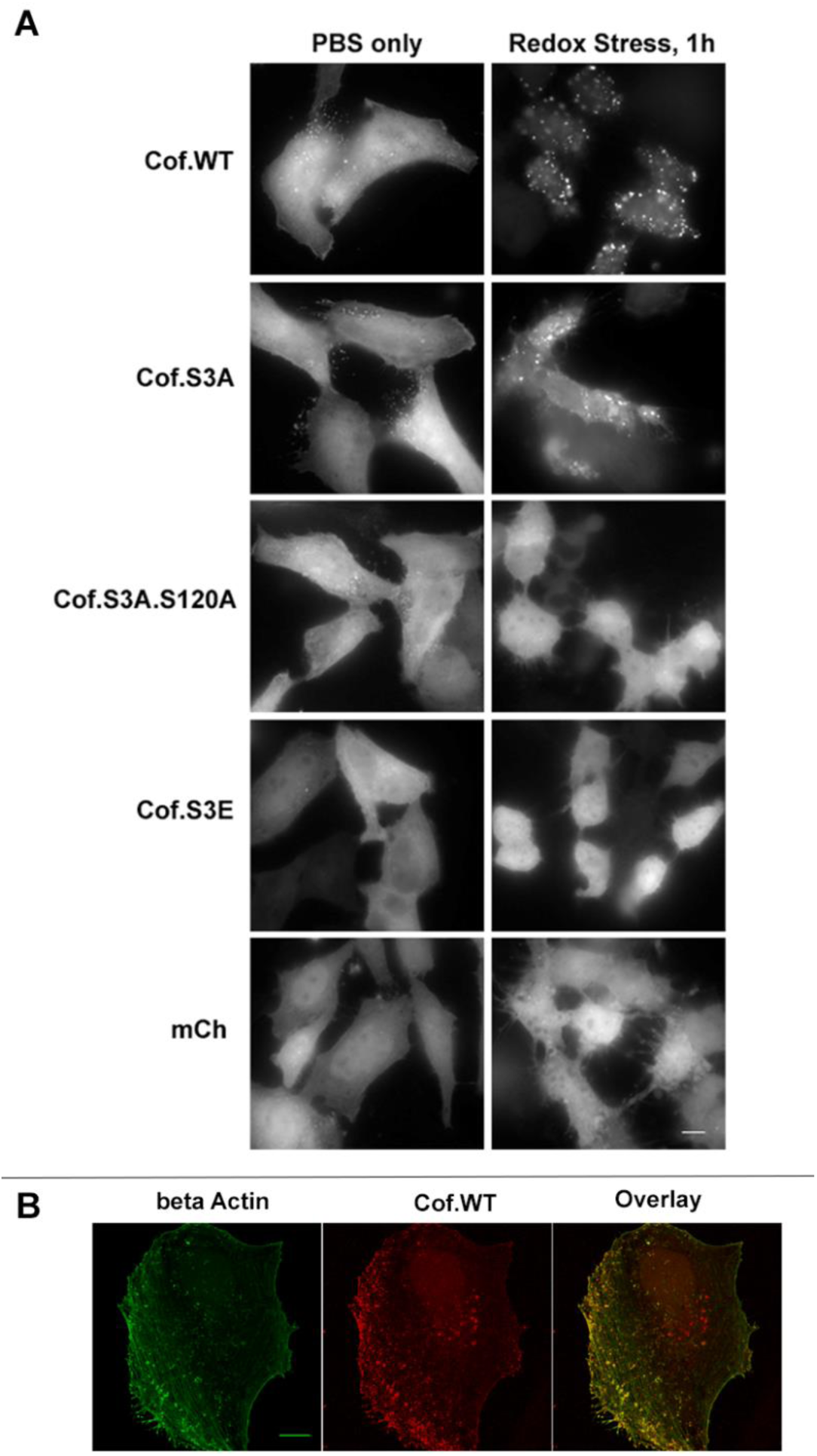
Cofilin Actin Rods Formed With CofilinWT Protein Fusion. **A.** Widefield fluorescent microscopy images of responses of cofilin protein fusions (expressed as a Cof-Cry2-mCherry fusion in HeLa cells) to oxidative and energetic stress. Actin binding cofilins (WT and S3A) incorporate into cofilin-actin rods after 1 hour in 6 mM D-deoxyglucose and 10 mM sodium azide, while actin-binding impaired mutants (S3A-A120A; S3E) and the cofilin-free control (Cry2-mCh) remain cytosolic. Scale bar = 10 microns. **B.** Zeiss Airyscan Confocal image of immunostaining of CofWT-Cry2-mCh cofilin-actin rods with beta Actin antibody demonstrates presence of endogenous actin in actin-cofilin clusters (yellow spots in overlay). Scale bars = 10 microns.

**FIGURE 2.**
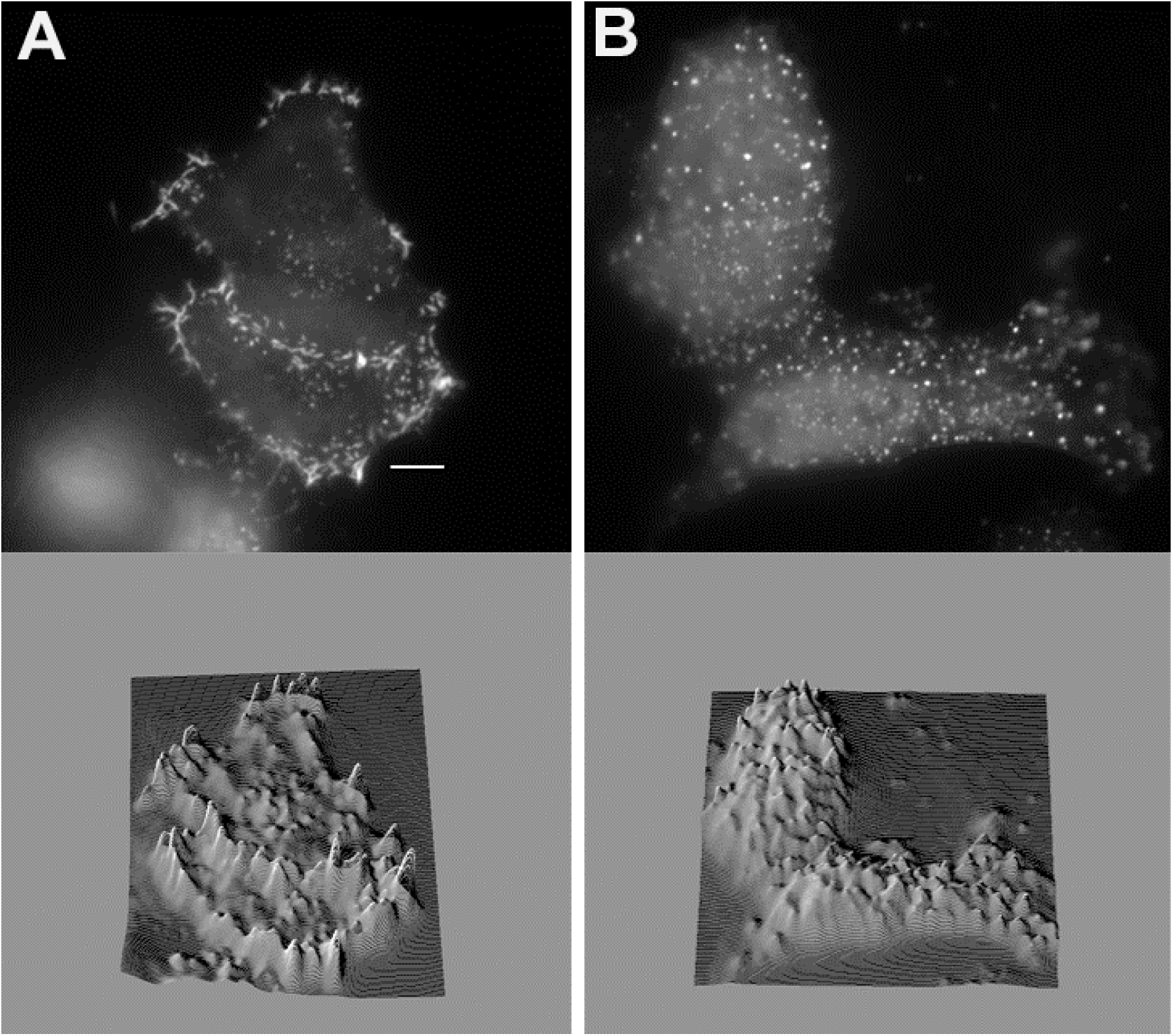
Response to variation in redox buffers. HeLa cells expressing Cof.WT-Cry2-mCh subjected to **A.** 6 mM D-deoxyglucose alone for 15 min at 37°C or **B.** 6 mM D-deoxyglucose/10 mM sodium azide for 15 min at 37°C. Bottom images depict 3D surface plots for each image (FIJI). Scale bar = 10 microns.

Having established that the cofilin-Cry2 fusions responded as expected to cofilin-actin bundle forming conditions (i.e., ATP depletion), we investigated whether the fusions would respond to both oxidative stress and light inputs in the presence of the Cry2 binding partner CIB. To test this, both N- and C-terminal cofilin-Cry2 fusion proteins and a β-Actin.CIB.GFP protein fusion were co-expressed in HeLa cells. Cells were incubated in the presence of ATP depletion medium for 15 min prior to imaging on a widefield fluorescent microscope. Of the various combinations of cofilin.Cry2 and actin.CIB, the N-terminal Cof.S3A.S120A and the C-terminal Cof.S3E exhibited robust light and redox dependent cofilin-actin fusion protein clustering that was reversible in the absence of blue light (Table I). As the response of the Cry2.mCh.CofS3E fusion was somewhat more robust (Figure 3 and **Supporting Movie I**), this construct was carried forward in conjunction with the β-actin.CIB.GFP construct for the remaining studies.

**FIGURE 3.**
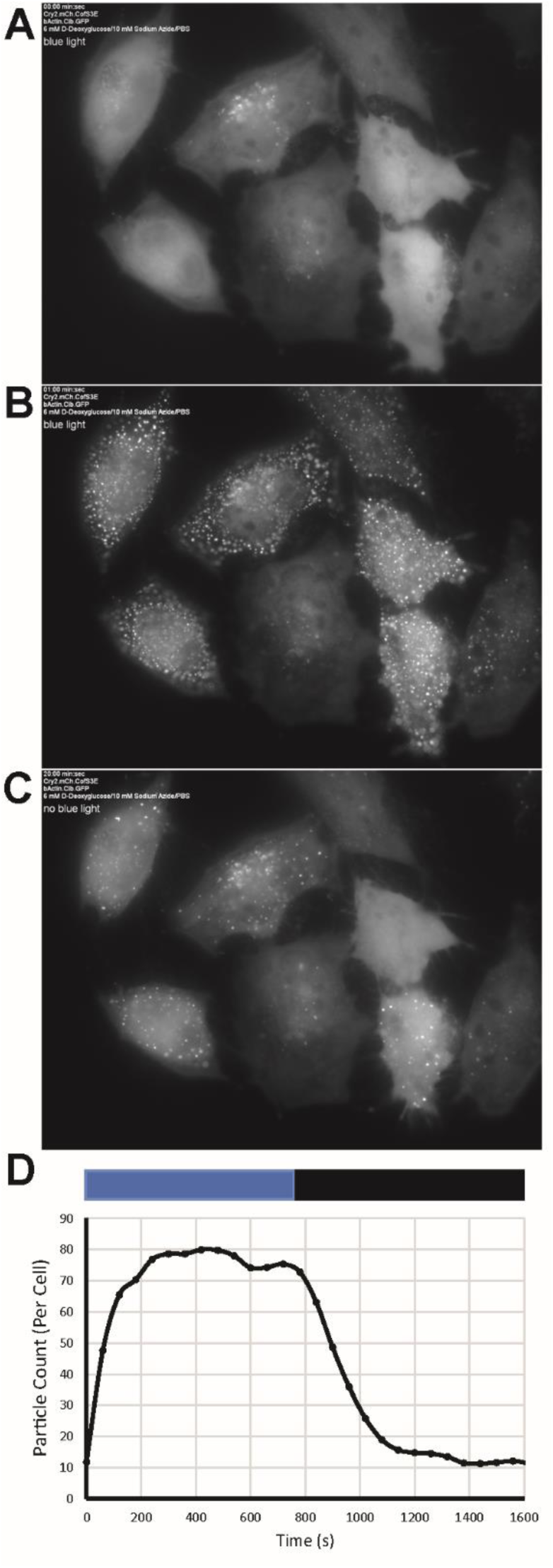
Demonstration of light initiated clustering between Cofilin and Actin linked Cry2/CIB. Light-initiated cofilin-actin clusters in HeLa cells form in the presence of oxidative and energetic stress. Prior to 470 nm light pulse (**A**) Cry2-mCh-CofS3E reagent is primarily cytosolic. Rods/clusters are clearly visible 1 min post-10 ms light pulse (**B**). 20 min post-pulse (**C**) most of the rods have dissipated. Line graph (**D**) shows automated particle counting analysis (FIJI) of cluster formation timecourse in cells from Supporting Movie I (blue bar = 470 nm on; black bar = 470 nm off).

**Table I.** SCREENING OF OPTOGENETIC CONSTRUCTS IN PRESENCE OF LIGHT + ATP DEPLETION. **Survey of light-stimulated Cofilin-Actin cluster formation**. Various N- and C-terminal cofilin mutant fusions to Cry2 were tested in the presence of betaActin.Cib.GFP and 10 mM Azide/6 mM D-deoxyglucose. Symbols: + + + + = >90% with light inducible clusters;+++ = ≥ 60 – 90% form clusters; ++ = 30 - 60% cells form clusters; + ≤ 30% cells form clusters -- = no cluster formation detected; I = irreversible cluster formation in the absence of blue light; R = reversible cluster formation in the absence of blue light.

| *Protein Pair* | *DMEM/10% FBS* | *ATP Depletion Media* |
| --- | --- | --- |
| ***N-terminal Cofilins*** |  |  |
| *Cof.WT-Cry2/bActin-Cib* | No clusters | ++++; I |
| *Cof.S3A.S120A-Cry2/bActin-Cib* | No clusters | +++; R |
| *Cof.S3E-Cry2/bActin-Cib* | No clusters | ++; R |
| *Cof.S3A.S120A-Cry2/Cib-ONLY* | No clusters | -- |
| ***C-terminal Cofilins*** |  |  |
| *Cry2-Cof.WT/bActin-Cib* | No clusters | ++++; I |
| *Cry2-Cof.S3A.S120A/bActin-Cib* | No clusters | +++; I |
| *Cry2-Cof.S3E/bActin-Cib* | No clusters | ++++; R |
| ***No Cofilin Controls*** |  |  |
| *Cry2-mCh/bActin-Cib* | No clusters | +; R |
| *Cry2-mCh/Cib-ONLY* | No clusters | -- |

As the interaction of the WT cofilin construct with endogenous actin varied in response to changing oxidative and energetic stress levels, the same treatments were applied to the optogenetic cofilin-actin bundling proteins (Figure 4; **Supporting Movies 2, 3, 4, and 5**). In response to a full complement of azide, but no glycolysis inhibitor (10 mM Azide/0 mM DDG), light activated cofilin-actin bundles were formed throughout cell bodies but were shorter lived than those formed in the presence of 10 mM Azide/6 mM DDG. In response to full inhibition of glycolysis, but no azide (0 mM Azide/6 mM DDG), light activated cofilin actin bundles formed chiefly around the cell periphery but were also shorter lived than those under full ATP depletion conditions. We note that this is an analogous response to that observed with incorporation of WT cofilin into endogenous actin bundles (Fig. 2). Finally, while control cells (in the absence of ATP depletion) form sparse light + oxidant induced clusters (Fig. 4B), weak light-activated localization between cytosolic Cry2.mCh.CofS3E and β-actin.CIB.GFP can still be observed in a subset of transfected cells (14.5% (10.7); n = 5 replicates; Supporting Figure 5 and **Supporting Movie 5**).

**FIGURE 4.**
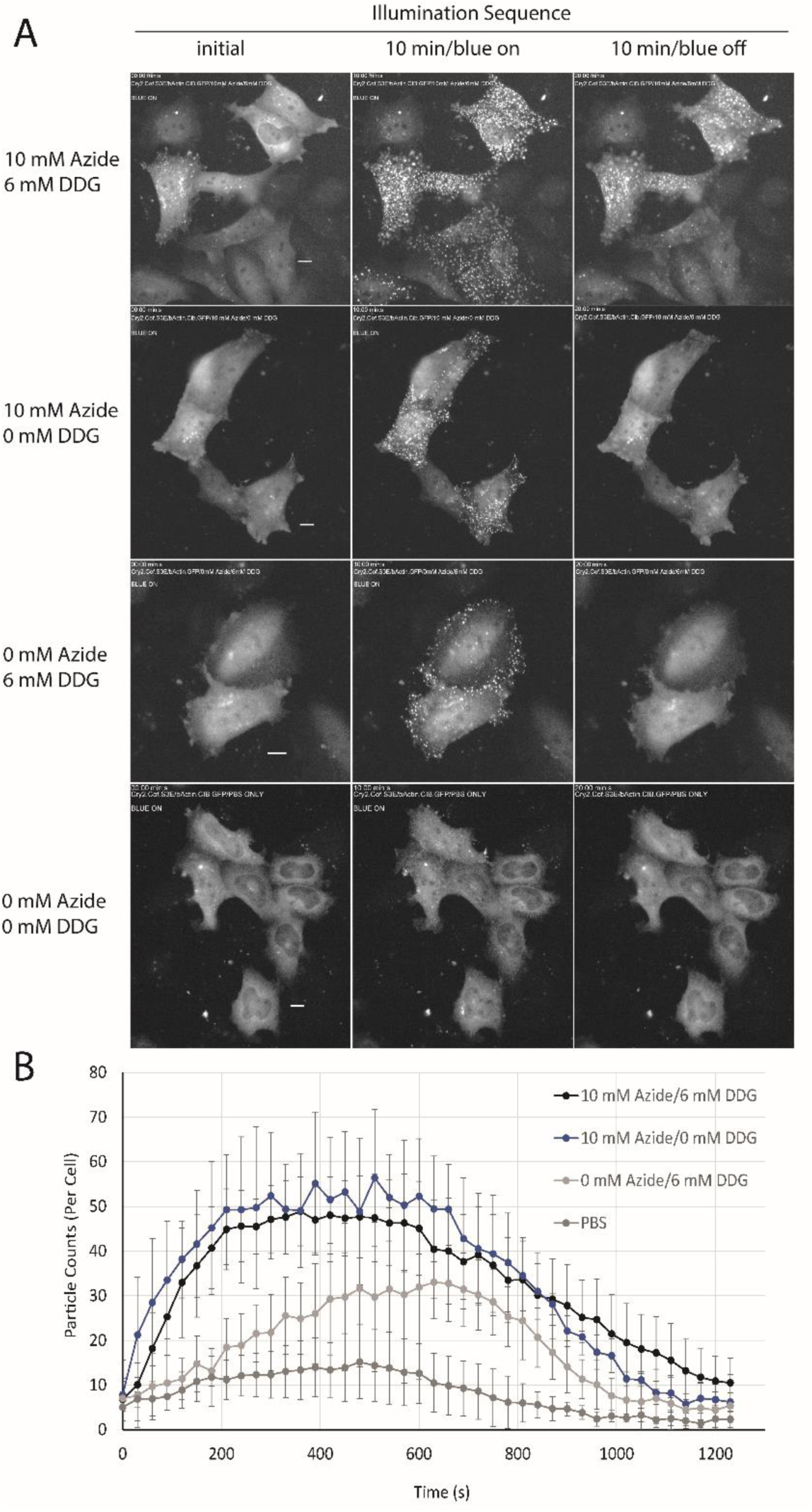
Light clustering behavior under variable oxidative stress conditions in HeLa cells. A. Images shown are pre-, 10 min post-, and an additional 10 min in the absence of blue light stimulation. Oxidative stress conditions are indicated in the figure for each group of cells; conditions applied for 10 min prior to beginning of illumination sequence. B. Counting of cofilin-actin clusters (ImageJ Particle Analysis) depicts timecourse of cluster formation in blue light (0 – 600 s) and dissipation in the absence of blue light (600 – 1200 s); Error bars represent SEM from n=3 replicate measurements. See also Supporting Movies 2 – 5.

The light and redox dependent clustering of the cofilin and actin fusions indicated that site directed mutagenesis of both cofilin, and actin might provide insight into the mechanism of interaction between the two proteins. One of the primary driving forces of native cofilin-actin bundling is the switch from primarily ATP-actin to ADP-actin that occurs as a result of oxidative stress-induced depletion of ATP. Thus, we investigated whether alteration of the ATP-binding site of actin via site directed mutagenesis might alter its ability to bind cofilin in the absence of oxidative stress by mimicking actin in its ADP bound state. Inspection of a crystal structure of actin with a non-hydrolysable ATP mimic (RCSB PDB 1NWK (37)) revealed two potential contacts that might be critical for binding the tertiary phosphate group of ATP: Ser14 (a predicted hydrogen bond contact) and Val 159 (potentially critical for maintaining binding cavity shape and hydrophobic packing) (Figure 5).

**FIGURE 5.**
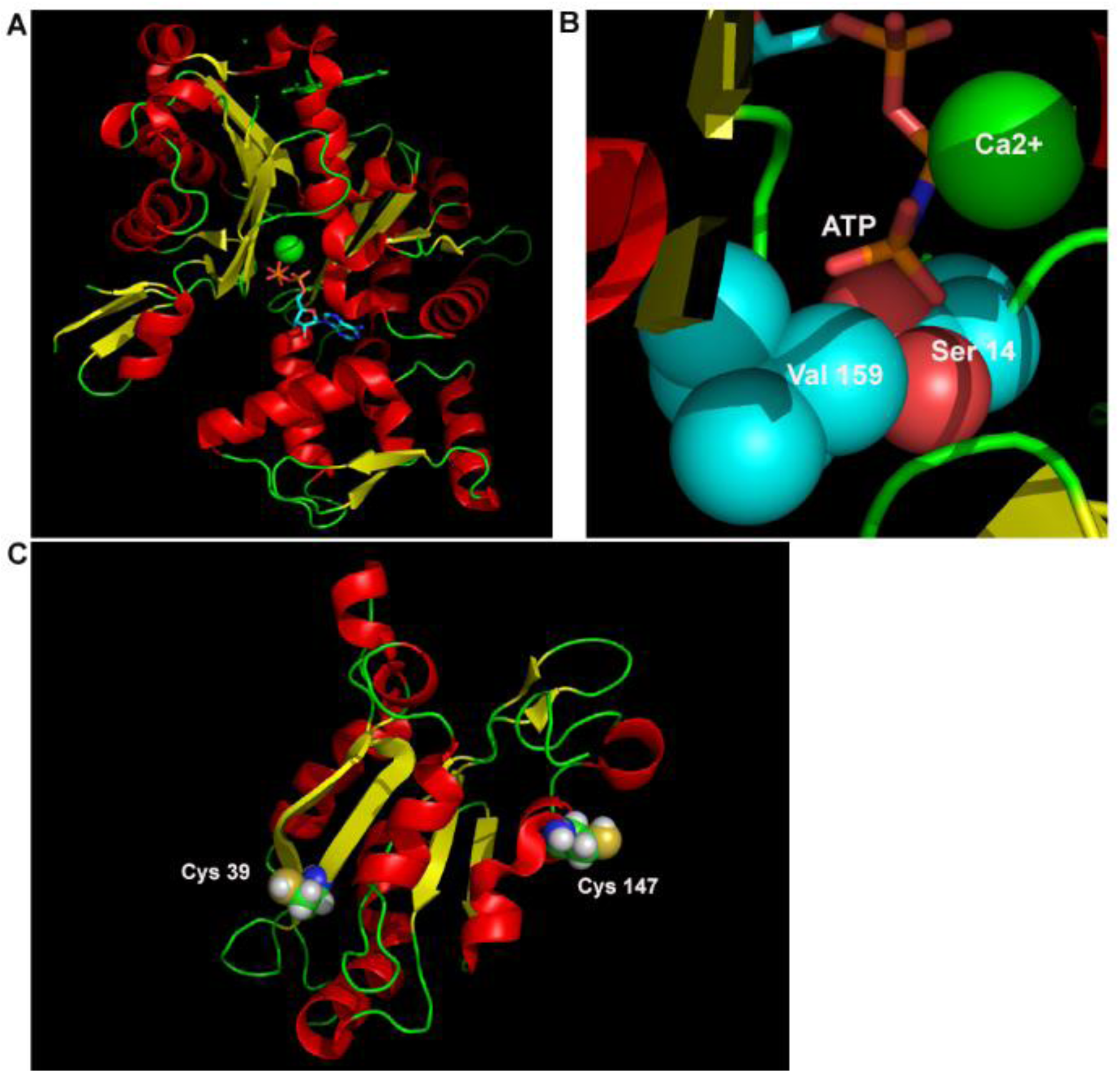
Mutagenesis strategy for modifying switch response. **A.** Crystal structure of monomeric Beta-actin bound to ATP analogue (RCSB PDB ID 1NWK) and **B.** a close up of the ATP binding pocket. The terminal phosphate of ATP is surrounded by a hydrophobic valine (Val 159) and hydrogen bound to a serine residue (Ser 14) and a calcium ion. Image rendered with PyMol. **C.** Cofilin cysteines (Cys 39; Cys 147; depicted as spheres color coded by element) responsible for intermolecular oxidation of Cofilin (RCSB PDB ID 1Q8G). Images rendered in PyMol.

We proposed that removal of the S14-phosphate hydrogen bond forming ability might create an actin mutant with the properties of an ADP-actin, whereas increasing the steric bulk of the Val 159 residue might have a similar effect by occluding the binding site for the tertiary phosphate of ATP. In addition to the proposed actin mutants, we also considered whether further mutagenesis of cofilin could impact the function of the light activated switch. Cofilin has four cysteine residues, two of which have previously been demonstrated to be critical for cofilin-actin rod stability via the formation of intermolecular disulfide bonds (Cys 39 and Cys 147; Fig. 5C (38)) (35, 39). Simultaneous mutation of cofilin cysteines 39 and 147 to alanine has been shown to interfere with efficient incorporation into cofilin-actin rods (35). We hypothesized that if cofilin-actin bundling with CofActor was similar to native cofilin-actin rod formation, then mutagenesis of cofilin Cys 39 and Cys 147 in CofActor should also interfere with switch function in the presence of oxidative stress.

Single and double mutant actins and cofilins (Table II) were generated and tested for their ability to participate in light and redox activated clustering when co-transfected with the corresponding Cry2.mCh.CofS3E or β-actin.CIB.GFP (Table II). In the case of the Val159Ile single mutant (β-actinV159I.CIB.GFP), light + redox stress were both required for cofilin-actin cluster formation. However, the Ser14Val mutant (β-actinS14V.CIB.GFP) gave a robust, reversible light activated clustering response in the absence of ATP depletion media (Figure 6 and **Supporting Movie 6**). Notably, the S14V mutant formed rod-like bundles with dimensions similar to those observed with native cofilin-actin rods (avg. 2 rods per cell (±1); length of 7.42 μm (±.68)). These structures are present prior to light activated recruitment of Cry2, indicating that they are formed independently of optogenetic cofilin. Optogenetic cofilin undergoes light-activated recruitment to these rod-like structures in addition to forming cofilin-actin clusters analogous to those observed in the presence of non-mutant actin (Fig. 6). Double mutants (S14V.V159L and S14V.V159I) exhibited an oxidative stress-independent response analogous to that of the S14V mutant, forming numerous light-independent rod-like structures present in cells (Fig. 6D). Interestingly, while the Val159Ile single mutant (β-actinV159I.CIB.GFP) did not efficiently form oxidative stress-independent clusters as observed with the S14V mutant, it did exhibit enhanced nuclear localization and formed numerous nuclear-localized light-activated cofilin-actin clusters in the presence of oxidative stress (Supporting Figure 6 and **Supporting Movie 7**), indicating that the presence of a hydrophobic sidechain at the 159 position may enhance nuclear import of the actin construct. In the case of the cofilin mutants (Figure 7), light and redox activation of cofilin-actin bundles were not significantly reduced in the C39A and C147A single mutants, but were highly diminished in the C39A.C147A double mutant, consistent with a previous report (35). Conversely, light + redox activation of bundle formation was eliminated in the C39D and C39D.C147D mutants, but not in the C147D mutant (Figure 7G).

**FIGURE 6.**
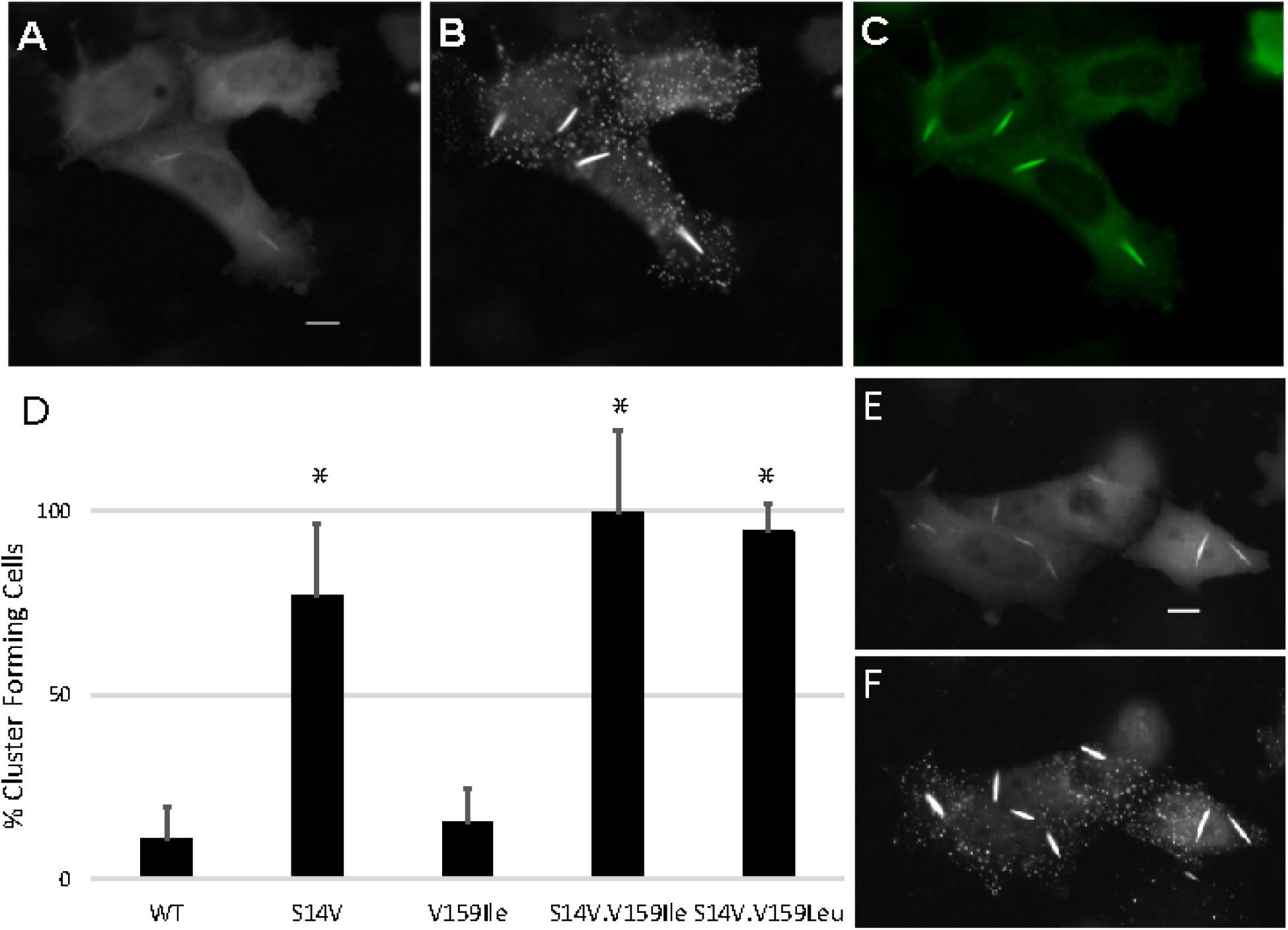
Rod formation with actin mutants in the absence of oxidative stress. **A.** HeLa cells transfected with Beta-actin.S14V.CIB.GFP/Cry2.mCh.Cof.S3E protein pair before (mCherry fluorescence shown) and **B**. 5 min post (mCherry fluorescence shown) blue light stimulation. **C.** Beta-actin.S14V.CIB.GFP construct at beginning of illumination sequence. **D**. % cells forming light inducible cofilin-actin rods and clusters in the absence of oxidative stress (n = 5 replicates per mutant; *P<0.001 versus WT control (One-way Anova; Holm-Sidak method)). **E**. Actin double mutant S14V.V159L.CIB.GFP/Cry2.mCh.Cof.S3E before and **F**. 10 min-post blue light illumination. All experiments were conducted under oxidant-free (PBS only) conditions. Scale bar = 10 microns. See also Supporting Movie 6.

**FIGURE 7.**
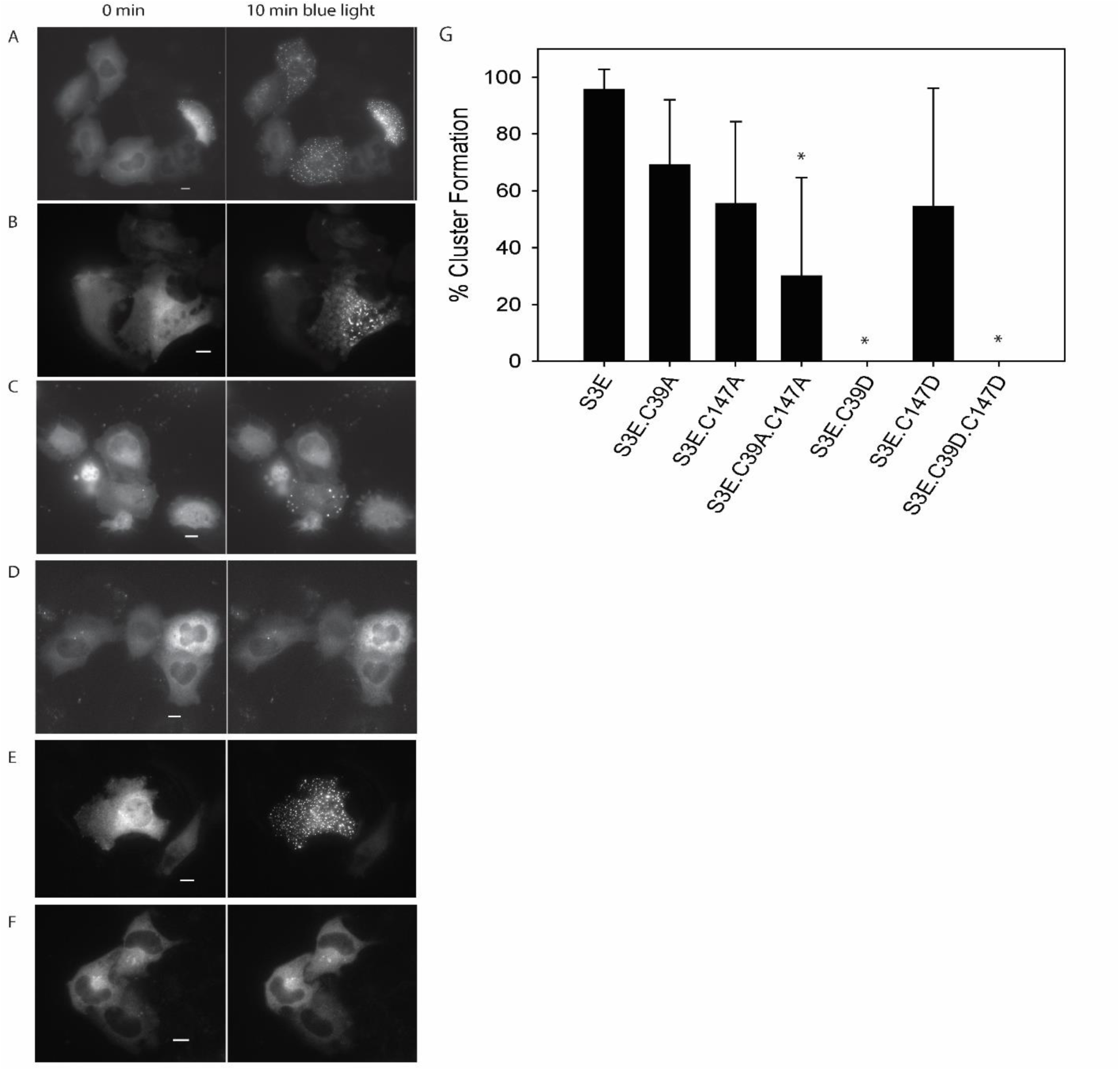
Mutation of Cofilin C39 and C147 impairs actin-cofilin bundle formation. Cells are shown pre- and post- 10 min of blue light stimulation. A. HeLa cells expressing Cof.S3E.C39A/Actin.CIB.GFP. B. HeLa cells expressing Cof.S3E.C147A/Actin.CIB.GFP. C. HeLa cells expressing Cof.S3E.C39A.C147A/Actin.CIB.GFP. D. HeLa cells expressing Cof.S3E.C39D/Actin.CIB.GFP. E. HeLa cells expressing Cof.S3E.C147D/Actin.CIB.GFP. F. HeLa cells expressing Cof.S3E.C39D.C147D/Actin.CIB.GFP. G. Quantification of A-F as % cells forming light inducible clusters; error bars are standard deviation from minimum 6 replicate measurements; (*P<0.001 versus S3E control; One-way ANOVA; Holm-Sidak method).

**Table II.** Summary of blue light activation responses of Actin and Cofilin Mutants under normal and oxidative stress conditions. Summary of light-stimulated cofilin-actin cluster formation of actin and cofilin mutants in the presence of 10 mM Azide/6 mM D-deoxyglucose/PBS or PBS. Symbols: + + + + = >90% with light inducible clusters;+++=50 – 90% form clusters; + = <50% cells form clusters; -- = no cluster formation detected.

| **ACTIN MUTANTS** | **PBS** | **ATP DEPLETED** |
| --- | --- | --- |
| *V159I* | -- | ++++ |
| *S14V* | +++ | ++++ |
| *V159L/ S14V* | ++++ | ++++ |
| *V159I/ S14V* | ++++ | ++++ |
| **COFILIN MUTANTS** |  |  |
| *S3E/C39A* | -- | +++ |
| *S3E/C147A* | -- | +++ |
| *S3E/C39A/ C147A* | -- | + |
| *S3E/C39D* | -- | -- |
| *S3E/C147D* | -- | +++ |
| *S3E/C39D/ C147D* | -- | -- |

Next, to demonstrate the potential utility of CofActor in neurons, we tested switch responsivity in primary hippocampal neuron cultures prepared from newborn mice. These cultures are routinely used as *in vitro* models to study cellular mechanisms of neurodegeneration (40). Prior to light stimulation, dissociated neuron cultures transfected with the CofActor system exhibited 8.14 (±2.49) actin rods and 2.14 (±0.55) cofilin rods in soma and 54.86 (±16.17) actin rods and 25.14 (±4.69) cofilin rods in neuronal processes (Figure 8). Upon stimulation with light in ATP depletion medium, the number of rods significantly increased in both subcellular compartments (Figure 8B). In the soma, the number of actin rods increased by 267% (21.71±5.32; p= 0.018), while the number of cofilin rods increased by 327% (7±2.02; p= 0.032). In neuronal processes, the number of actin rods increased by 168% (92.43±15.7; p= 0.004), while the number of cofilin rods increased by 223% (56.14±12.73; p= 0.039). Taken together, the data presented here shows that the CofActor optogenetic sensor reports on conditions required for the formation of native actin-cofilin rods in cells exposed to ATP depletion conditions and therefore is a promising tool to study the cellular mechanisms that lead to abnormal cytoskeletal reorganization and are associated with pathological conditions, including neurodegeneration and excitotoxicity.

**FIGURE 8.**
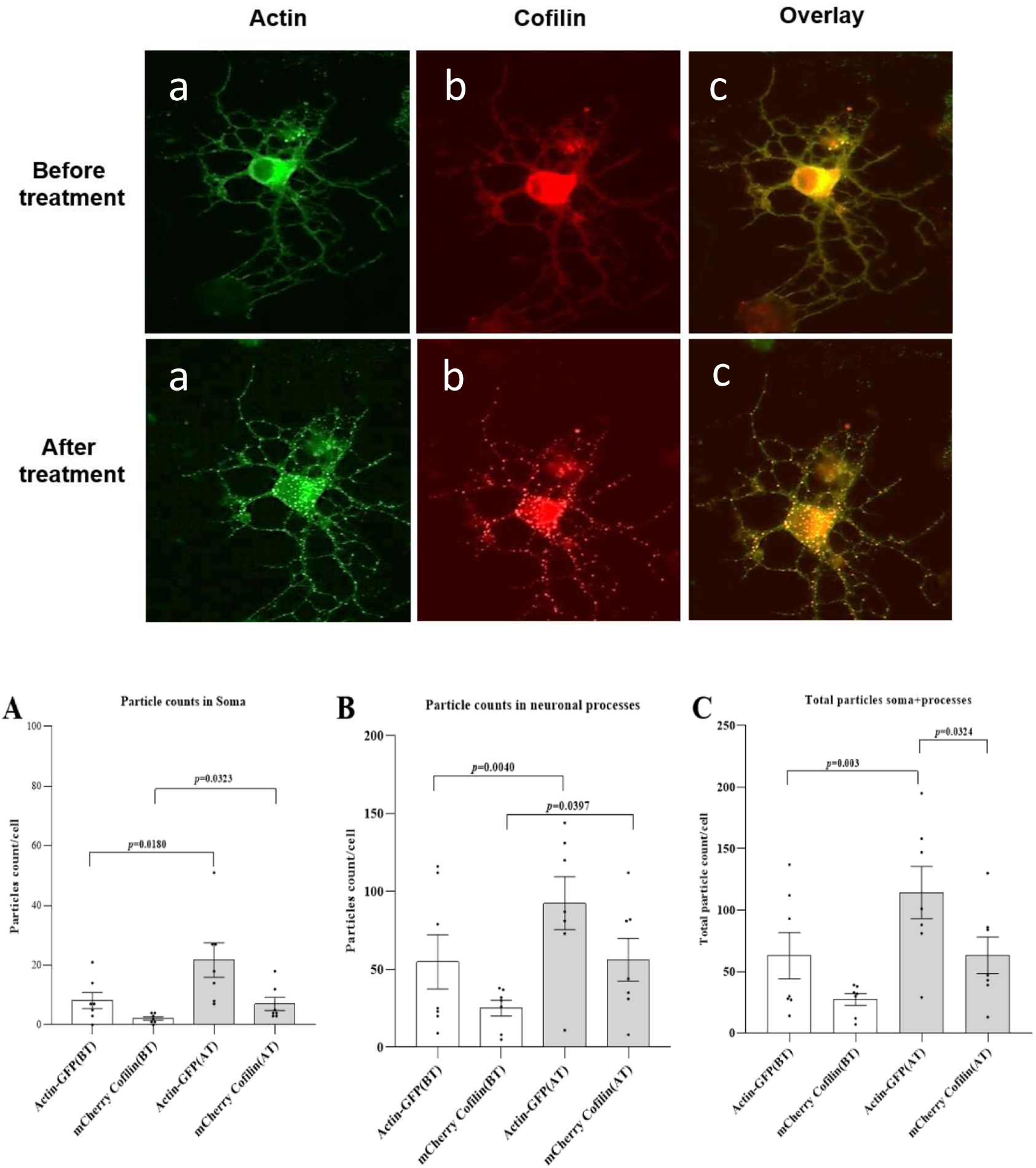
Validation of CofActor in primary hippocampal neuron cultures. **A**. Representative micrographs of primary hippocampal neurons transfected with Cofactor system, before (a, b, c) and after (a’, b’, c’) exposure to ATP depletion conditions and illumination sequence. **B**. Number of actin and cofilin rods increases significantly in neurons transfected with the Cofactor system upon ATP depletion and illumination with blue light both in soma (**A**) and in neuronal processes (**B**). Graph **C** shows the overall increase in rod formation in neurons. Data is average of 7 neurons from 2 independent cultures. Error bars are S.E.M.

## DISCUSSION

Cytoskeletal dysregulation is a prominent feature of numerous diseases, including cancer, stroke, and neurodegenerative disorders (19, 41, 42). Whether prominent cytoskeletal features are progenitors, or merely symptoms of disease progression remains open for debate (43). As a result, biosensors and bioactuators that recapitulate the anomalous cytoskeletal responses to disease are valuable for contextualizing their role(s) in disease pathology. In this work, a known, disease relevant protein-protein interaction (cofilin/actin) has been coupled with an optogenetic oligomerizer (Cry2/CIB), to better understand the factors governing their interaction(“CofActor”). Where previous studies have developed mutants of cofilin that resist incorporation into cofilin-actin rods under normal physiological conditions (30), these mutants resist incorporation until activated with a pulse of blue light. This is advantageous for cellular studies, as cells can be more precisely treated with redox stimuli, and the cytoskeletal events are precisely triggered following application of the stress stimuli. Furthermore, the two-component nature of the Cry2/CIB system enables testing of both cofilin and actin mutants in parallel, whereas a cofilin-only or actin-only system only allows testing with endogenous actin or cofilin. The S14V actin mutant, which eliminates the need for ATP depletion medium, likely operates by shifting actin into either an ADP-bound state or promotes an actin structure similar to the ADP-bound state. This is supported by the numerous rod-like actin substructures that form from the actinS14V.CIB.GFP construct which are similar to cofilin-actin rods formed *in vivo*. Perhaps significantly, these rod-like actin structures are present prior to CofActor activation with blue light. These pre-assembled rods may incorporate significant amounts of endogenous cofilin, whereas the absence of endogenous cofilin in these rods would indicate an interesting actin-nucleated mechanism for rod formation that is independent of cofilin. Whether rods are able to form *in vivo* in a cofilin independent fashion is unknown, and invites further inquiry. While the S14V single and double actin mutants exhibit robust light activated clustering in oxidant-free medium, they also function in the presence of ATP depletion buffer, making them potentially useful redox-independent light activated tools that function under various experimental conditions (Table II). We expect that the mutagenesis of other critical contacts between ATP and actin may also generate interesting redox-independent behaviors when incorporated into the CofActor system. Alternately, site directed mutagenesis of cysteine residues in cofilin, guided by previously reported studies (35, 36), resulted in Cry2PHR.mCh.Cofilin fusions with impaired light activated cluster formation. We were skeptical as to whether cysteine mutagenesis would produce a significant effect, given that Cry2PHR.mCh.CofS3E readily formed light-induced clusters with the ActinS14V mutant under oxidant-free conditions. Nonetheless, the C147A.C39A cofilin double mutant significantly reduced the light-induced clustering behavior in the presence of ATP depletion medium. The effect is notably more pronounced with Cys to Asp than with Cys to Ala mutations. In particular, the Cys39Asp mutation is sufficient for eliminating light activated clustering altogether under oxidative stress. This effect may result from electrostatic repulsion between the Asp sidechain and the actin binding interface when brought into close proximity with actin, as has been proposed elsewhere(36). Alternately, the Cys to Ala mutants can compensate somewhat for the loss of the native disulfide bond capability through non-specific hydrophobic interactions with the binding interface and are thus less deleterious to cofilin-actin cluster formation.

Several possible applications are currently under investigation for CofActor. For example, the CofActor switch could form the basis of a Boolean AND gate for induction of protein-protein interactions, where both light and redox stress inputs would be required for clustering.

CofActor could also be forming the basis for an “optical two-hybrid” assay for actin – actin binding protein interactions, with cluster formation as a readout of protein-protein interaction propensity rather than transcription of a reporter gene. We anticipate that the capability to produce cofilin-actin bundles with photostimulation could be a useful tool in the field of neuroscience to study the effects of abnormal cytoskeletal re-arrangement associated with synaptic dysfunctions and behavioral deficits in neurological disease model organisms.

## Authors’ contributions

RMH lab: RMH designed experiments. FBS, ABK, and RMH performed cloning and cell culture experiments. FBS, ABK, and RMH analyzed data and prepared figures. EMS lab: EMS and RMH designed experiments. EMS, VVP and WPB prepared neuron cultures and performed neuronal experiments. VVP and WPB analyzed the data and prepared figures. All authors (FBS, VVP, ABK, WPB, AM, EMS, and RMH) contributed to writing the manuscript and agreed to the content of this paper.

## Acknowledgements

RMH lab: Authors wish to thank Elizabeth Ables and Karen Litwa for assistance with Zeiss Airyscan imaging. EMS lab: Authors wish to thank to Rachel Hope Dodson for outstanding technical support and Heather Sorenson for research coordination support.

## SUPPORTING MOVIES

**Supporting Movie 1**: Light activation of HeLa cells expressing Cry2PHR.mCh.CofS3E/Actin.CIB.GFP treated with 10 mM NaN3 and 6 mM D-Deoxyglucose.

**Supporting Movie 2**: Light activation of HeLa cells expressing Cry2PHR.mCh.CofS3E/Actin.CIB.GFP treated with 10 mM NaN3 and 6 mM D-Deoxyglucose.

**Supporting Movie 3**: Light activation of HeLa cells expressing Cry2PHR.mCh.CofS3E/Actin.CIB.GFP treated with 10 mM NaN3 and 0 mM D-Deoxyglucose.

**Supporting Movie 4:** Light activation of HeLa cells expressing Cry2PHR.mCh.CofS3E/Actin.CIB.GFP treated with 0 mM NaN3 and 6 mM D-Deoxyglucose.

**Supporting Movie 5:** Light activation of HeLa cells expressing Cry2PHR.mCh.CofS3E/Actin.CIB.GFP treated with 0 mM NaN3 and 0 mM D-Deoxyglucose.

**Supporting Movie 6:** Light activation of HeLa cells expressing Cry2PHR.mCh.CofS3E/ActinS14V.CIB.GFP in DPBS.

**Supporting Movie 7:** Light activation of HeLa cells expressing Cry2PHR.mCh.CofS3E/ActinV159Ile.CIB.GFP treated with 10 mM NaN3 and 6 mM D-Deoxyglucose.

---

**Supporting Figure 1.**
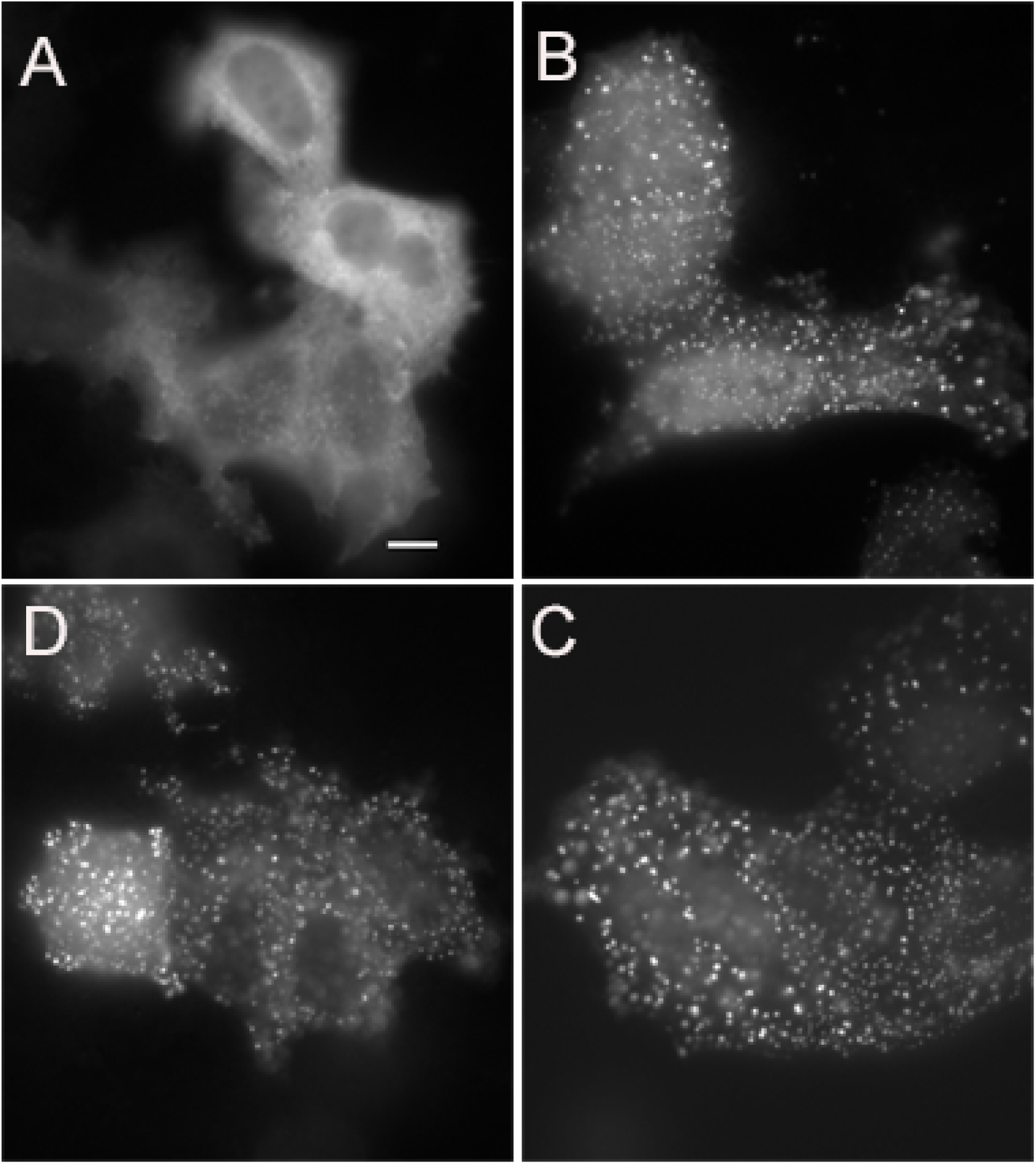
Time course of CofWT.Cry2.mCh/ β-actin clustering in HeLa cells incubated in 10 mM NaN_3_/6 mM D-deoxyglucose/DPBS. **A)** no treatment (PBS only); **B)** 15 min post-treatment; **C)** 30 min post-treatment; **D)** 45 min post-treatment. Images acquired with confocal microscope (60X objective; scale bar = 10 microns).

**Supporting Figure 2.**
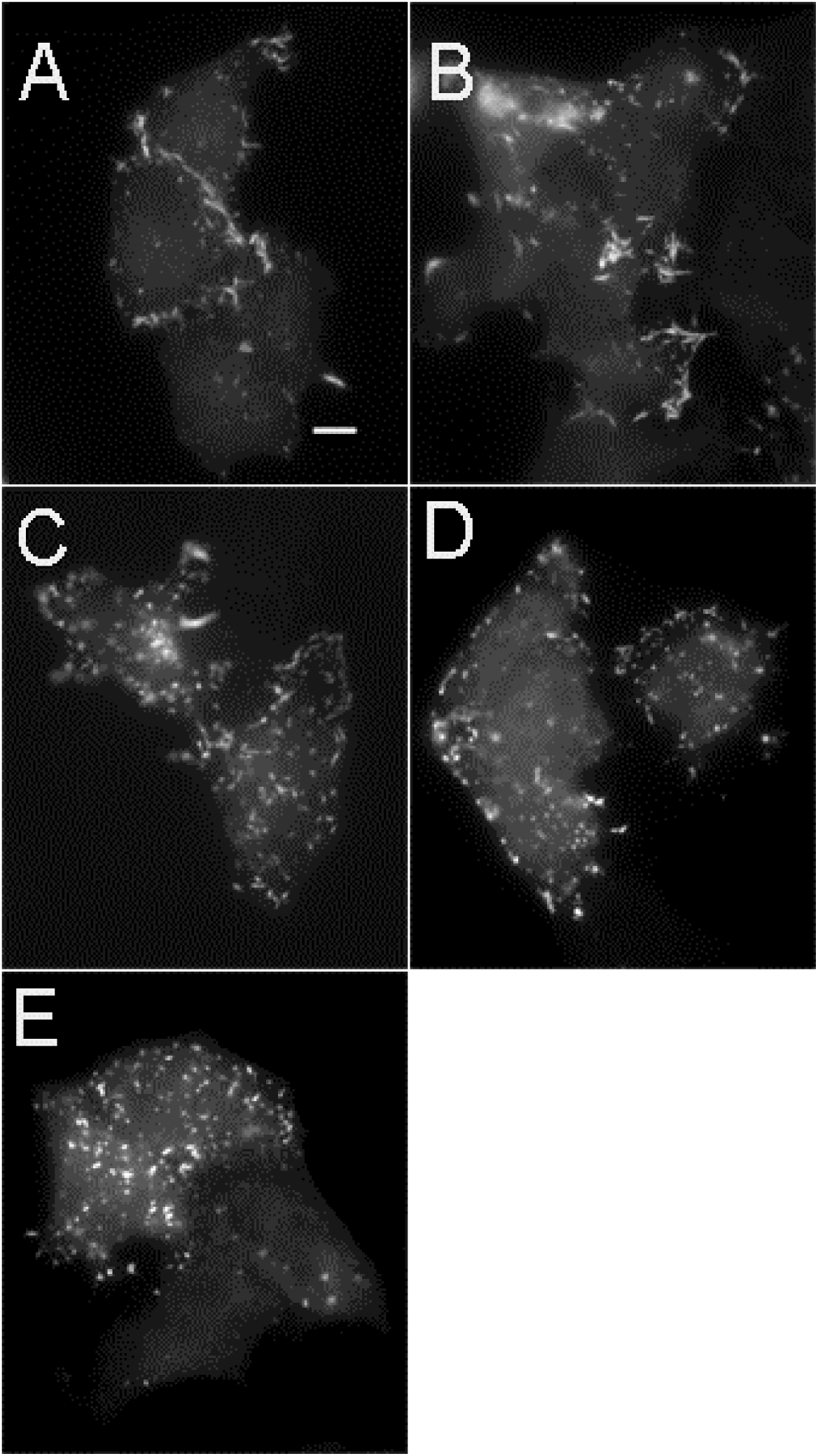
CofWT.Cry2.mCh/β-actin clustering in HeLa cells incubated with increasing concentrations of NaN3 and 6 mM D-deoxyglucose/DPBS for 15 min. **A)** 0 M NaN_3_; **B)** 50 μM NaN_3_; **C)** 100 μM NaN_3_; **D)** 200 μM NaN_3_ **E)** 400 μM NaN_3_. Images acquired with confocal microscope (60X objective; scale bar = 10 microns).

**Supporting Figure 3.**
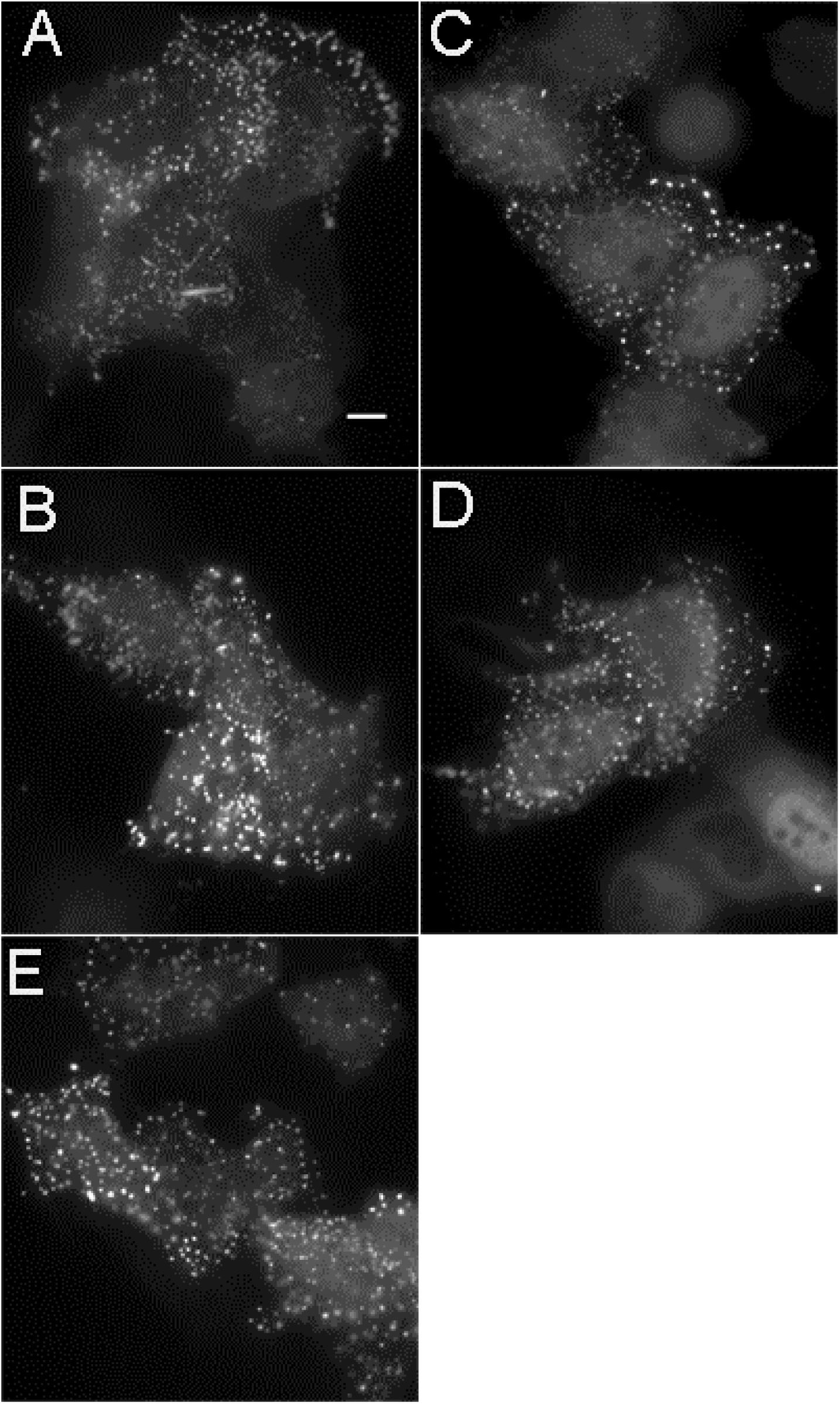
CofWT.Cry2.mCh/β-actin clustering in HeLa cells incubated with increasing concentrations of DDG and 10 mM azide/DPBS for 15 min. **A)** 0mM **B)** 1mM **C)** 2mM **D)** 3mM **E)** 5mM. Images acquired with confocal microscope (60X objective, scale bar = 10 microns).

**Supporting Figure 4.**
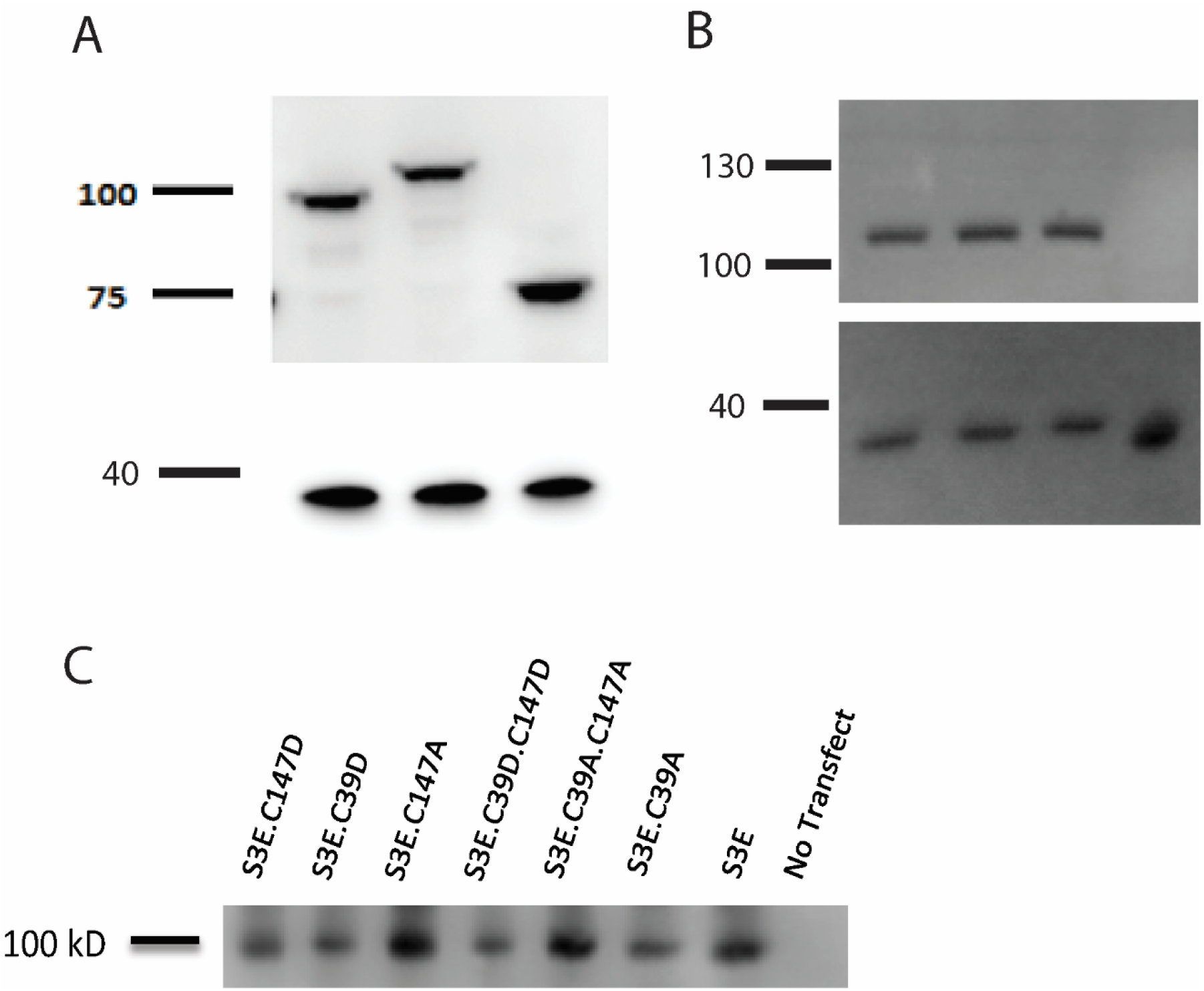
**A.** anti-mCherry Western blots of (L – R) C- and N-terminal cofilin-Cry2 fusions (Cry2.mCh.Cof; Cof.Cry2.mCh) vs. Cry2.mCh control. GAPDH loading control appears at the expected MW of approx..37 kD. **B.** anti-GFP Western blots of (L – R) βactin.S14V.CIB. GFP, βactin.V159Ile.CIB.GFP; actin, βactin.S14V.V159Leu.CIB.GFP, and an untransfected control lysate. GAPDH loading control appears at the expected MW of approx..37 kD. **C.** anti-mCherry Western blots of Cry2.mCh.Cof mutant proteins.

**Supporting Figure 5.**
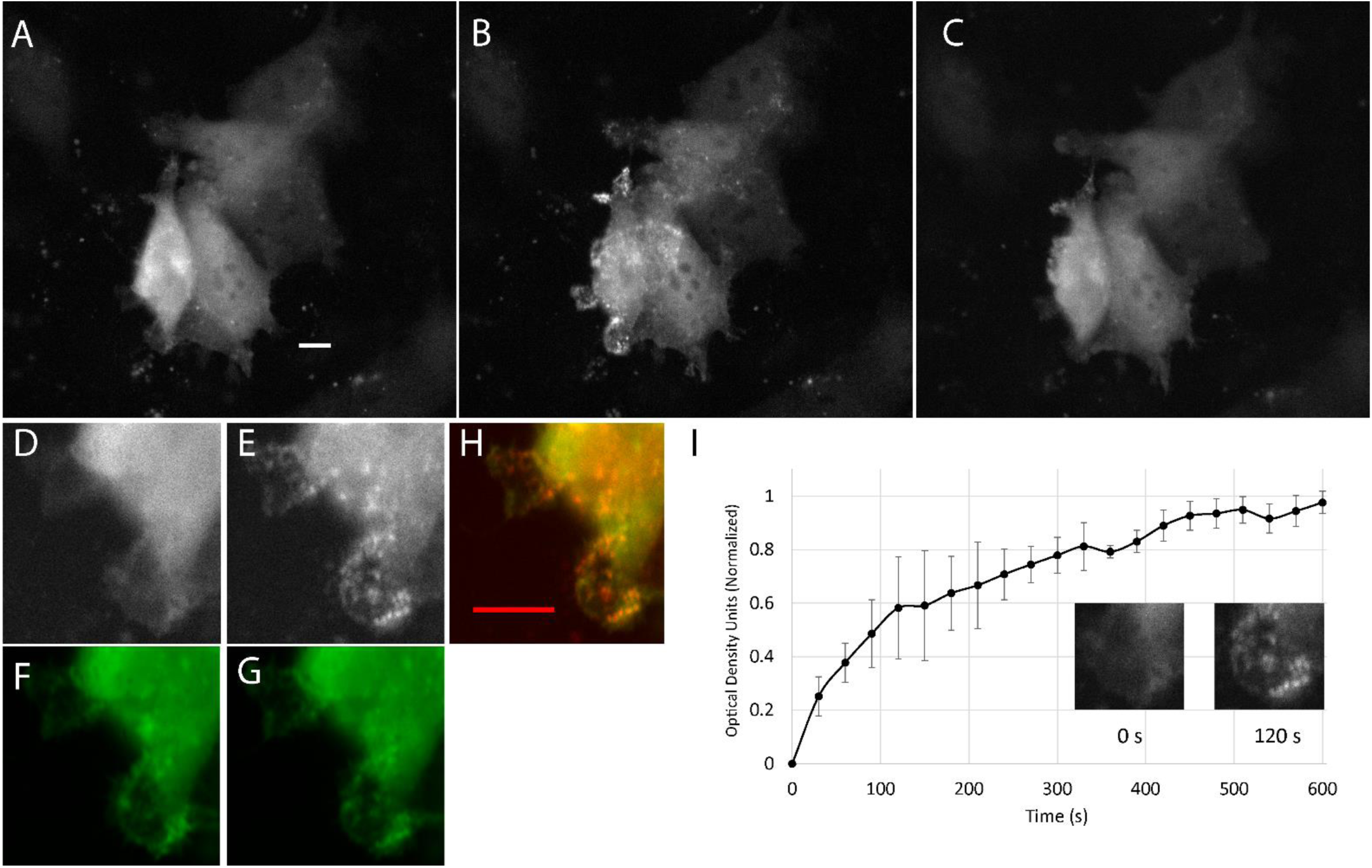
Redox-independent co-localization of Cry2.mCh.Cof.S3E and bActin.CIB.GFP at the cell periphery. HeLa cells expressing Cry2.mCh.Cof.S3E/Actin.CIB.GFP are shown **A.** pre-, **B.** post - 10 min of blue light stimulation, and **C.** after an additional 10 min without blue light stimulation. Zoomed region of cell from 5A, shown cell periphery (**D.** mcherry; **F.** GFP) before and (**E.** mcherry; **G.** GFP) 2 min post-light stimulation. **H.** Co-localization of images shown in E. and G. Scale bar = 10 microns. **I.** Timecourse of co-localization of cropped region shows change in mCherry fluorescence intensity vs. time at the cell periphery. Error bars (SEM) are determined from three different peripheral regions.

**Supporting Figure 6.**
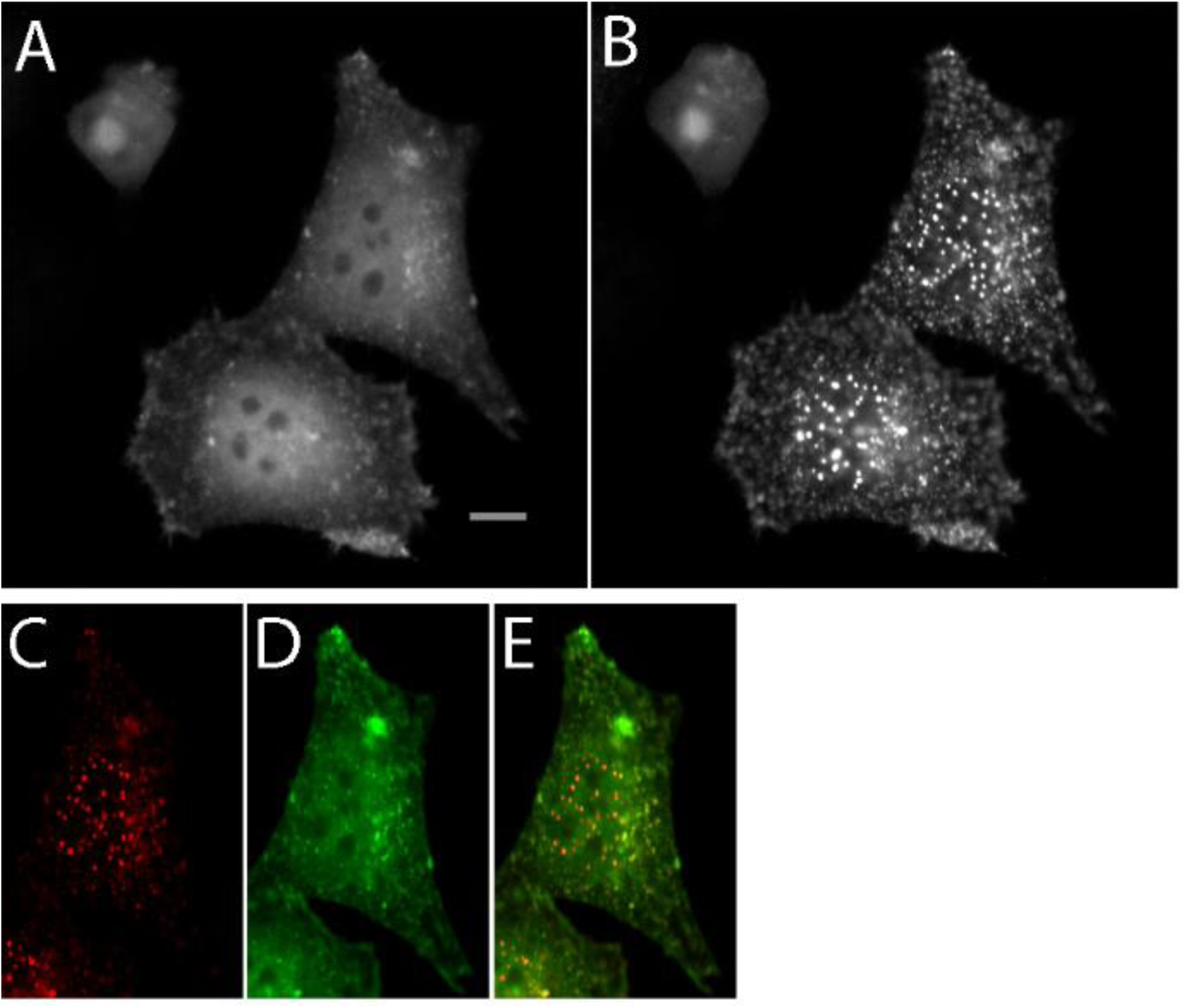
Light inducible nuclear cofilin-actin cluster formation. HeLa cells expressing Cry2.mCh.Cof.S3E /Actin.V159Ile.CIB.GFP in 10 mM azide/6 mM DDG/PBS are shown **A.** pre- and **B.** post - 5 min of blue light stimulation. Nuclear-localized cofilin-actin clusters shown in **C.** Cry2.mCh.Cof.S3E (mCherry fluorescence), **D.** Actin.V159Ile.CIB.GFP (GFP fluorescence), and **E.** overlay of mCherry and GFP channels. Scale bar = 10 microns. See also Supporting Movie 7.

